# A gene set collection of RBP target genes

**DOI:** 10.1101/2025.10.14.682362

**Authors:** Markus Glaß, Stefan Hüttelmaier

## Abstract

RNA binding proteins (RBPs) are key post-transcriptional regulators controlling every aspect of the RNA life cycle from synthesis to decay. We extracted RBP target genes from publicly available enhanced cross-linking and immunoprecipitation followed by sequencing (eCLIP-seq) data of 168 RBPs and assembled a gene set collection that can be used to examine gene lists for enriched RBP targets via functional enrichment analysis methods like over-representation analysis (ORA), gene set enrichment analysis (GSEA) or gene set variation analysis (GSVA).

## 1 Background & Summary

RNA binding proteins (RBPs) comprise a class of proteins that bind to and interact with RNA molecules. The interactions between RBP and RNA can affect all aspects of the RNA’s life cycle, including synthesis, processing, post-transcriptional modification, localization, stability and degradation, in sum positioning RBPs as key regulators of RNA fate.

Functional enrichment analysis (FEA), also referred to as gene set analysis (GSA) methods encompass computational approaches used to determine whether certain properties of gene products appear more frequently than expected by chance. These properties can be multifaceted, e.g., the the involvement in a particular biological pathway or disease, common chromosomal location or common targets of a regulatory molecule like an RBP. Several distinct FEA methods have been developed, including over-representation analysis (ORA), gene set enrichment analysis (GSEA) and gene set variation analysis (GSVA). For ORA, selected genes are tested whether certain properties are over-represented among these genes. A background list of detected genes is essential for proper comparison to the list of selected genes [1].

GSEA takes a different approach by considering the entire ranked list of genes (usually ranked by fold change between two experimental conditions) rather than just a subset. It calculates an enrichment score based on where genes from a particular gene set fall in the ranked list. This method does not require arbitrary cutoffs for defining significant genes [2]. GSVA, however, calculates enrichment scores for individual samples rather than comparing groups of samples. It uses a non-parametric approach that ranks genes within each sample and calculates sample-wise gene set enrichment scores as a function of genes inside and outside the gene set [3]. Thus, scores produced by GSVA can be interpreted as the degree to which a gene set is up- or down-regulated in each individual sample. Besides these three particularly popular methods, a multitude of further classes of FEA have been developed. An overview of methods and implementations can be found in [4].

Despite the differences in the underlying statistics, all FEA methods rely on gene sets that provide the assignments between genes and the properties of their products. The Molecular Signatures Database (MSigDB) [5] represents a comprehensive resource of gene sets applicable to FEA methods. The Human MSigDB is divided into nine collections, each representing gene sets focusing on different aspects, like the hallmarks, representing well-defined biological states or processes [6] or the curated collection that contains gene sets derived from research papers and databases. Gene sets can be obtained from the MSigDB website (https://www.gsea-msigdb.org) as gene set (sub-)collections, stored in gmt format, a tab-delimited text file format, containing one gene set per line. While the MSigDB also contains a collection of regulatory target gene sets, this collection is restricted to microRNA and transcription factor targets. Thus, although the curated collection encompasses gene sets containing target genes of a few selected RBPs, e.g., BRCA1, IGF2BP1 and IGF2BP2, a comprehensive and focused collection of RBP targets is missing.

In a large-scale effort, van Nostrand et al. [7] performed enhanced crosslinking and immunoprecipitation followed by sequencing (eCLIP-seq) binding assays for a multitude of RBPs in human cell lines and deposited the results at the ENCODE data portal [8] (www.encodeproject.org). These data exhibit a great resource of genome-wide binding sites of currently more than 150 distinct RBPs. These data allow the deduction of target transcripts as well as general binding characteristics like sequence context around binding sites by inspecting the supplied data for peak regions.

We gathered high-confidence eCLIP data of 168 distinct RBPs provided by the ENCODE data portal. From the obtained peak files, we extracted genes associated with significantly enriched peaks and assembled a gene set collection representing associations of these 168 RBPs and the genes their target transcripts originated from. The collection is stored in the gmt file format and can directly be used with stand-alone programs like GSEA [2] (www.gsea-msigdb.org/gsea/) or R-packages like clusterProfiler [9] and GSVA [3].

## 2 Methods

### 2.1 Data Collection

Relevant data sets were identified from the ENCODE data portal [8] (www.encodeproject.org) by selecting the “eCLIP” assay on the “Experiment Search” page. Subsequently, irreproducible discovery rate (IDR) filtered peak files, containing significantly enriched eCLIP peaks reproducible between two independent biological replicates were obtained from the respective project pages in bed file format. The full list of RBPs including ENCODE file identifiers is provided in table S1.

### 2.2 Data Processing

Genes covered by peaks were obtained by applying the intersect command from the bedtools suite (v2.31.1) [10] to obtain strand-specific overlaps between peaks and genes as annotated by ENSEMBL GRCh38 v114 [11]. From the resulting intersection files, gene names were extracted and a gmt file was created that contains one gene set for each RBP in each available cell line.

## 3 Data Record

The gene set collection is comprised of 243 gene sets containing putative target genes of 168 distinct RBPs determined from either hepatoblastoma-derived Hep-G2 or chronic myeloid leukemia-derived K-562 cells or both (Fig. 1A). In general, more RBP target genes were derived from eCLIP-data originating from Hep-G2 cells than from K-562 cells (median target gene number 860.5 vs. 596; Fig. 1B). Notably, Jaccard-indexes, measuring the similarity between gene lists extracted from eCLIP data, displayed rather low concordance between target genes extracted from Hep-G2 and corresponding K-562 data sets of the same RBPs (median Jaccard-index 0.21; Fig. 1C). This might reflect differences in the technical feasibility of eCLIP in dependence of the respective cell models as well as differences in the overall expression profiles of the distinct cell lines. Biological functions of the RBPs included in the gene set collection covered processes like splicing, gene expression regulation, translation and localization (Fig. 1D).

**Figure 1:**
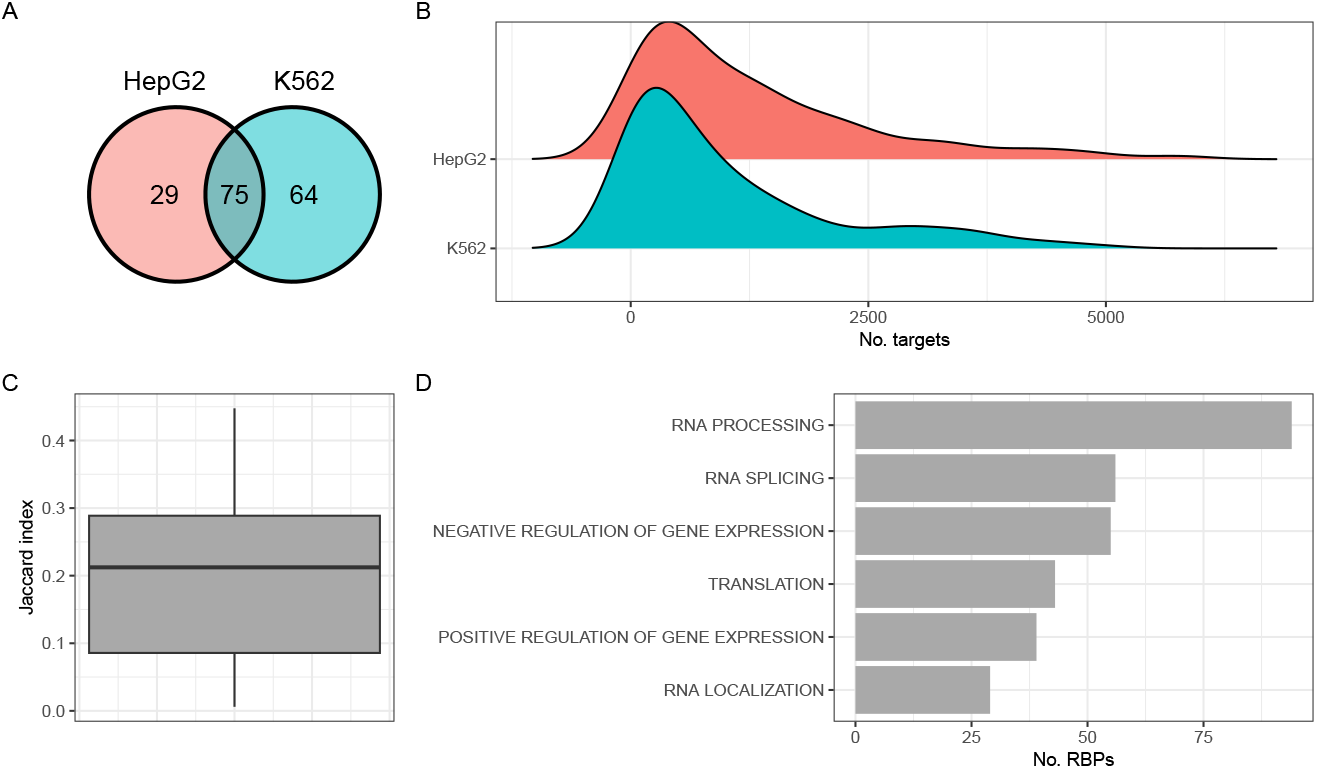
Overview of the RBP gene set collection. **A** Distribution of eCLIP experiments among the two applied cell lines Hep-G2 and K-562. **B** Distribution of derived target gene numbers among the two distinct cell lines. **C** Jaccard-indexes for those 75 eCLIP-experiments performed for the same RBP in both cell lines. **D** Selected Gene Ontology-derived biological processes in which the RBPs from the gene set collection are involved.

The gene set collection is available as gmt file at Zenodo (DOI 10.5281/zenodo.17296967) and from GitHub (github.com/HuettelmaierLab/RBP_Gene_Set_Collection). Each line of the gmt file comprises names of genes crosslinked to a specific RBP in either Hep-G2 or K-562 cells. The name field, i.e., the first entry of each line, specifies the RBP and cell line, the second field (description) represents the ENCODE identifier of the peak file of which the target genes were obtained from.

## 4 Technical Validation

To test usability and plausibility of our gene set collection, we used it in conjunction with different R-packages to perform ORA, GSEA and GSVA using publicly available gene lists as input.

For testing plausibility, we first performed an ORA using a gene list derived from a PAR-CLIP [12] experiment using endogenous Pumilio 2 (PUM2) protein for pull-down in colon cancer-derived HCT-116 cells. The gene list obtained from supplemental data of Lee et al. [13] contained 370 distinct genes. In contrast, the respective PUM2 eCLIP experiment, incorporated in our gene set collection, contained almost 3000 target genes. Of those, 198 were also contained in the PAR-CLIP list. Despite this large difference in target gene numbers, ORA results, obtained by using the enricher function of the clusterProfiler R-package [9], revealed PUM2 targets to be the most significantly enriched gene set, containing the most genes of the input PAR-CLIP list (Fig. 2A). Thus, PUM2 target genes derived from an orthogonal method could be identified as being enriched for PUM2 targets using our eCLIP-derived RBP target gene set collection.

**Figure 2:**
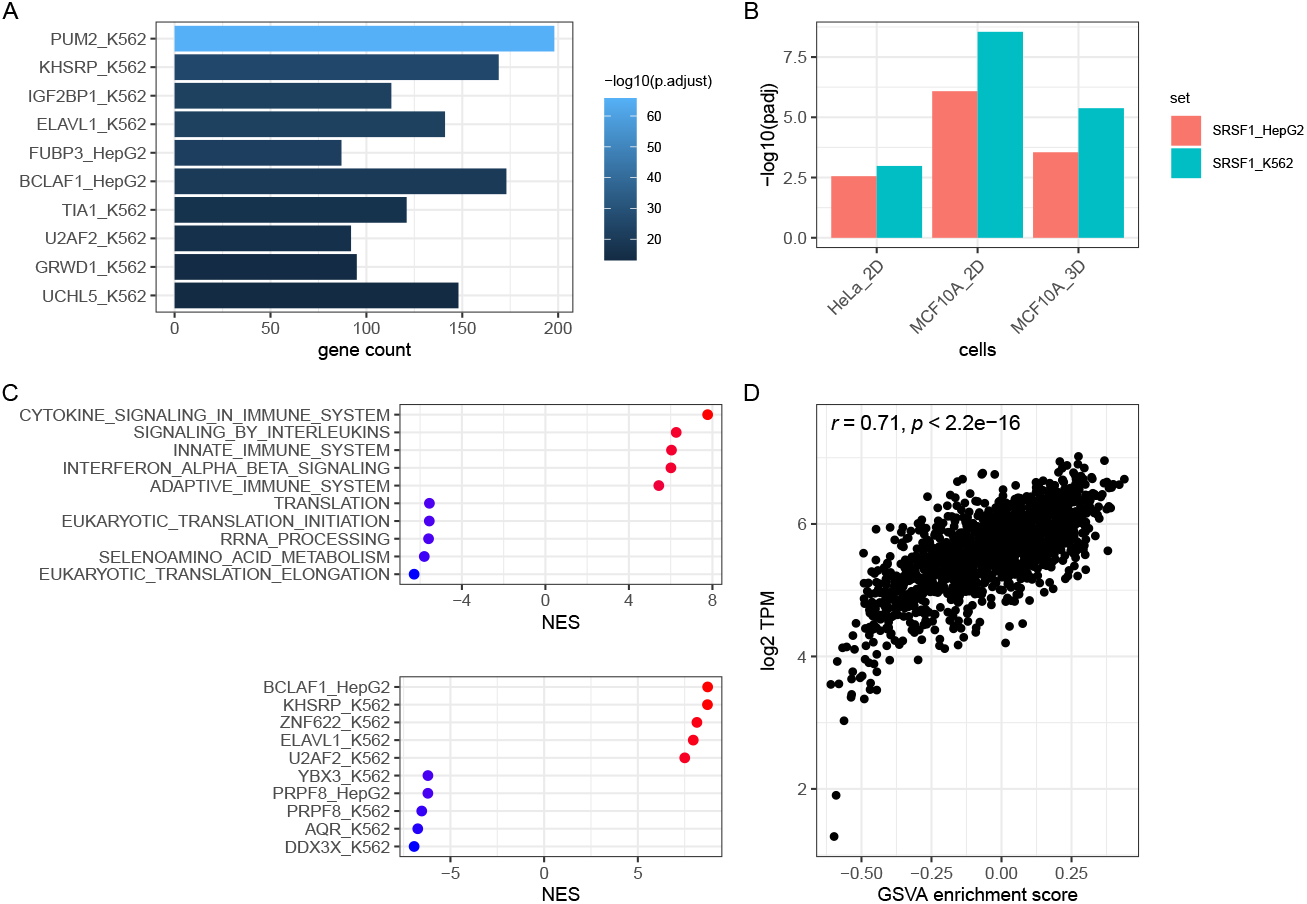
Functional enrichment analysis (FEA) results using the proposed RBP target gene set collection. **A** Top 10 most significantly enriched RBP gene sets using the PUM2 PAR-CLIP-derived genes as input gene list for an ORA. **B** ORA-derived FDR adjusted p-values regarding enrichment of SRSF1-related gene sets using lists of genes alternatively spliced upon SRSF1 overexpression in the indicated cell models as input. **C** Normalized enrichment scores (NES) of the top 5 positively and negatively enriched Reactome (upper panel) and RBP gene sets (lower panel) upon GSEA using log2 fold changes of gene expression upon cGAMP stimulation. All displayed gene sets were significantly enriched (FDR adjusted p-values < 0.05). **D** GSVA enrichment scores of ELAVL1 target genes and ELAVL1 RNA expression from 1404 distinct CCLE cell line data sets.

Since pure binding, as determined by methods like PAR-/eCLIP, not necessarily reflects functional consequences, we next tested if genes associated with functions of a certain RBP could be used to refer to this RBP as well. Therefore, we used lists of genes identified to be alternatively spliced upon over-expression of the splicing factor SRSF1 for ORAs. These gene lists were generated by analyzing RNA-seq data of control cells in comparison to cells over-expressing SRSF1 for alternative splicing events by Anczuk et al. Experiments were performed in three different cell models, 2D-cultured cervical cancer-derived HeLa cells, 2D-cultured breast cancer-derived MCF-10A cells as well as 3D-cultured MCF-10A cells [14]. SRSF1 target gene sets derived from Hep-G2 and K-562 cells were significantly enriched (FDR adjusted p-value ≤ 0.05) upon ORA using the detected alternatively spliced genes from all three cell models as input (Fig 2B). To test the gene set collection for applicability with GSEA, we obtained a list of gene expression fold changes between control and cyclic GMP-AMP (cGAMP)-stimulated monocytic THP-1 cells. cGAMP was used as a surrogate for viral infection by acting as agonist of the immune adaptor stimulator of interferon (STING). The authors of this study performed further high-throughput sequencing assays, including PAR-CLIP to conclude that upon cGAMP stimulation the RBP ELAVL1 is activated by post-translational modification leading to increased stabilization of its target transcripts [15]. GSEA performed using R/clusterProfiler [9] on the MSigDB curated sub-collection containing Reactome pathways [16], revealed several gene sets related to the innate immune response as being the strongest positively enriched gene sets. Gene sets showing the strongest negative enrichments were connected to mRNA translation (Fig 2C). GSEA performed with the RBP collection revealed targets of immune system-related RBPs, including ELAVL1 as being the most strongly positively enriched sets [17, 18, 19, 20]. Furthermore, the gene sets showing the strongest negative enrichment were associated with RBPs related to the immune system as well (DDX3X, AQR), but also associated with RBPs known to regulate RNA translation (DDX3X, YBX3) [21, 22, 23, 24]. GSEA using the stand-alone GSEA application [2] in pre-ranked mode and the classic scoring scheme yielded highly similar results (data not shown).

Finally, we tested our gene set collection in conjunction with GSVA. We obtained RNA expression values of 1404 distinct cell line models available provided by the cancer cell line encyclopedia via the R-package ExperimentHub [25, 26] and used this dataset as input for the gsva function of the GSVA R-package [3]. After obtaining enrichment scores of all RBP gene sets, we calculated the correlation between ELAVL1 RNA expression and GSVA enrichment scores of ELAVL1 target genes. This revealed a strong positive correlation suggesting that ELAVL1 exerts a promoting impact on its RNA targets supporting its reported stabilizing role [15] (Fig 2D).

## Data Availability

The most recent version of the gene set collection is available in gmt file format from github.com/HuettelmaierLab/RBP_Gene_Set_Collection. The version described in this manuscript is available at Zenodo (DOI 10.5281/zenodo.17296967).

## Code Availability

R-scripts and gene lists for performing all described FEAs are available at Zenodo (DOI 10.5281/zenodo.17296967).

## Author Contributions

MG: conceptualization, data curation, formal analysis, visualization and writing - original draft. SH: supervision and writing - review and editing.

## Additional Information

Supplementary information can be found in table S1.

## Competing Interests

The authors declare no competing interests.

## Acknowledgements

We thank the ENCODE project and the lab of Gene Yeo for providing publicly available eCLIP data via the ENCODE portal.

## Funding

This work was supported by funding of Deutsche Forschungsgemeinschaft (DFG) to SH (RU5433, P9:468534282) and MG (GRK2751, 449501615)

## Notes

https://github.com/HuettelmaierLab/RBP_Gene_Set_Collection

https://doi.org/10.5281/zenodo.17296967

### Competing Interest Statement

The authors have declared no competing interest.

